# Innovative airborne DNA approach for monitoring honey bee foraging and health

**DOI:** 10.1101/2025.04.17.649222

**Authors:** Mateus Pepinelli, Alejandro José Biganzoli-Rangel, Katherine Lunn, Patrick Arteaga, Daniel Borges, Amro Zayed, Elizabeth L. Clare

## Abstract

Environmental DNA refers to genetic material collected from the environment and not directly from an organism of interest. It is best known as a tool in aquatic ecology but eDNA has been found associated with almost every substrate examined including soils, surfaces, and riding around on other animals. The collection of airborne eDNA is one of the most recent advances used to monitor a variety of organisms, including plants, animals, and microorganisms. Evidence suggests a high turnover rate providing a recent signal for the presence of DNA associated with an organism. Here, we test whether biological material carried in air in honey bee colonies can be used to evaluate recent foraging and colony health. We sampled air using purpose built “bee safe” filters operating for 5-6 hours at each colony and successfully recovered plant, fungal and microbial DNA from the air within honey bee colonies over a 3-week pilot period. From these data we identified the core honey bee microbiome and plant interaction data representing foraging behaviour. We calculated beta diversity to estimate the effects of apiary sites and sampling date on data recovery. We observed that variance in ITS data was more influenced by sampling date. Given that honey bees are generalist pollinators our ability to detect temporal signals in associated plant sequence data suggest this method opens new avenues into the ecological analysis of short term foraging behavior at the colony level. In comparison variance in microbial 16S sequencing data was more influenced by sampling location. As the assessment of colony health needs to be localized, spatial variance in these data indicate this may be an important tool in detecting infection. This pilot study demonstrates that colony air filtration has strong potential for the rapid screening of honey bee health and for the study of bee behaviour

## Introduction

Honey bees (*Apis mellifera* Linnaeus 1758) play a crucial role in pollination and ecosystem functioning, and significantly impact plant reproductive success (Klein et al. 2007). Their conservation has become a critical area of focus in ecological and agricultural research due to their essential role in diverse ecosystems (Potts et at. 2016). They contribute to the pollination of approximately 75% of the fruits, vegetables, and nuts grown in the United States, making them integral to food security and biodiversity (Klein et al. 2007). However, honey bee populations have been declining due to multiple factors including habitat loss, pesticide exposure, climate change, and diseases. According to French et al (2024), honey bees grapple with an average of 23 stressors simultaneously, which translates into a large number of potential interactions, all with the capacity to undermine their health.

Environmental DNA is an important and well-established tool in ecology with applications in biodiversity surveys (Denier et al. 2017), community ecology (Garrett et al. 2023), monitoring (Mathieu et al. 2020), species interactions (Banerjee et al. 2022) and the detection of rare and invasive species (Thomas et al. 2020). It has been best established as a tool in aquatic ecology but is a growing method in health surveillance and monitoring of diseases, particularly those of ecological threat. For example, case studies of surveillance for fungal oak wilt diseases (Gautheir et al. 2023), pathogens in amphibians (Hall et al. 2015) and freshwater infectious parasites (Thomas et al. 2022) have been presented and a recent special issue of the journal Environmental DNA was devoted to eDNA in disease ecology (Childress et al. 2024).

The use of eDNA to document the interactions of colonial honey bees with their environment has a relatively long history in the eDNA literature. Honey has been the common source for eDNA, for example Giersch et al. (2009) used honey to detect the presence of the microsporidian parasite *Nosema ceranae*, one of the most widespread causes of colony disease, and Laube et al. (2010) developed primers to document plants visited by bees. This type of work has been greatly expanded over the last decade for the study of insect-plant interactions (Schnell et al. 2010), hive foraging (e.g. Hawkins et al. 2015, de Vere et al. 2017) and general hive health (Ribani et al. 2020). Other approaches using molecular tools to document foraging and hive health have included the use of hive swabbing and debris (Boardman et al. 2024) and commercial hive health diagnostics of adults, generally requiring 100-300 individuals from a brood frame (e.g. National Bee Diagnostic Centre, Alberta, Canada, https://www.nwpolytech.ca/nbdc/) for PCR testing. It has been debated which of these techniques are, strictly speaking, environmental DNA. For example, microbial or pollen metabarcoding (Bell et al. 2017, Wizenberg et al. 2024) may not strictly qualify as eDNA but is often included when part of general environmental sampling, making the distinction largely academic when collection and analysis overlap. One challenge with honey and debris as eDNA sources is determining the temporal scale of the signature. Honey is not deposited in a particular sequence within the hive and honey cells may carry over from year to year. As a consequence, unless the capping of specific cells is being monitored, determining the timing of DNA signatures in honey is likely impossible making it hard to track the temporal patterns of foraging. Swabbing individuals for eDNA and using pollen-traps for pollen metabarcoding (Wizenberg et al. 2023, Wizenberg et al. submitted) does track recent foraging but may be invasive, require lethal sampling and disrupt beekeeping daily activities.

As an alternative to determine the temporal variability in foraging, a common eDNA approach is to look for insect DNA deposited directly on flowers (Gomez et al. 2023). This approach, colloquially called “shake and soak”, involves a large number of flowers pooled into one water based soaking procedure (Wari et al. 2024). The bottle of flowers is then shaken up to try to dislodge insect DNA. The water is filtered using a standard aquatic eDNA filter from which the DNA can be extracted. This procedure can provide much finer resolution on foraging but has three main challenges. First, while using the pooling approach can increase throughput and might be necessary to increase PCR success, it may also lead to the masking of DNA templates or rare species (Drinkwater et al. 2021). Second, the insect visitor cannot be determined ahead of time so it is more efficient as a generalist approach to floral visitation rather than to study a particular pollinator. Finally, it is becoming established that insect DNA travels through air (Roger et al. 2022) just like plant (Johnson et al. 2019) and animal (Clare et al. 2022, Garrett et al. 2023) eDNA, and it is therefore likely that at least some of these insect signatures may not reflect foraging but simply an accumulation of local biodiversity. In fact, washing leaves for insect DNA is an established method of general screening for biodiversity (Valentin et al. 2020, Macher et al. 2023), thus it may be complicated to distinguish pollination from general insect presence in the environment.

One alternative for tracking the health and ecology of colonial honey bees is through sampling of the colony air for biological material. Airborne eDNA sampling for biodiversity is an emerging tool that utilizes genetic material suspended in the air to detect and monitor biodiversity (Johnson et al. 2019, Clare et al. 2021). This approach allows for the collection of DNA from a variety of organisms, including plants, animals, and microorganisms, providing a comprehensive snapshot of the local ecosystem (Littlefair et al. 2023), but it can also be targeted when an organism is frequently contained in an enclosed or semi-enclosed space (Clare et al. 2021, Garrett et al. 2023). Airborne sampling for pathogens has a much longer history (Whon et al. 2012, Jiang et al. 2015) than sampling for plants and animals, but the technique is rapidly expanding for terrestrial monitoring. For our purpose we thus consider all samples collected using an eDNA technique to be a mix of eDNA and cellular (e.g. microbe) sources and treat them as one sample. The advantage of air as an eDNA source for tracking the ecology of honey bees is that it likely provides a very recent eDNA signal. Current data on the duration of airborne signals suggest they last only a matter of hours (Garrett et al. 2023) before they start to degrade and thus provide a solution to the indeterminate timing of ecological information from honey and colony debris.

Here we use filtered colony air from honey bee hives to determine whether we can simultaneously assess colony health and foraging ecology in one non-invasive approach. As global concerns about pollinator decline escalate, there is a growing need for efficient and sensitive monitoring methods (Potts et al., 2010) which provide information on a narrow time period of activity by the target organism. Airborne eDNA has been advocated as a tool to enhance our ability to study pollination systems (Keller et al. 2015, Clare et al. 2021), which are vital for ecosystem functioning and agricultural productivity. In this study we test the potential of airborne sampling of honey bee hives for two purposes; 1) to determine whether air in hives can provide insights into recent foraging of bees and 2) to determine whether air samples can be used to measure microbial health of colonial bees.

## Materials and Methods

### Sampler design

To collect air from inside hives we used a modified version of the 12V mini samplers described by Garrett et al. 2023 (Design 2). In brief, these consisted of a 12 V, 0.15A 1.8W WINSINN 40 mm × 40 mm × 20 mm DC brushless fan modified with a 2 pin adapter and connected to a 12V “car charger” type power source. These were powered by RoyPow 30 W PD Power Banks (RoyPow USA), which have a compatible 12V “cigarette” lighter socket. Each fan has a 3D printed frame held in place using nuts and bolts, and we modified the top clamp to be a ‘BeeSafe” red version with a top grid so the bees could not land on the filter (Figure 1). The filter itself was a sterilized square cut from a Filtrete 1900 Smart Air Filter designed for standard home furnace filtration. We disassembled the furnace filter under UV lights in a laminar flow cabinet, cut sampling squares, and then UV sterilized these prior to use following Garrett et al. (2023). A video on sampler construction and assembly is provided at https://youtu.be/DHHR5Cg5dUE.

**Figure 1:**
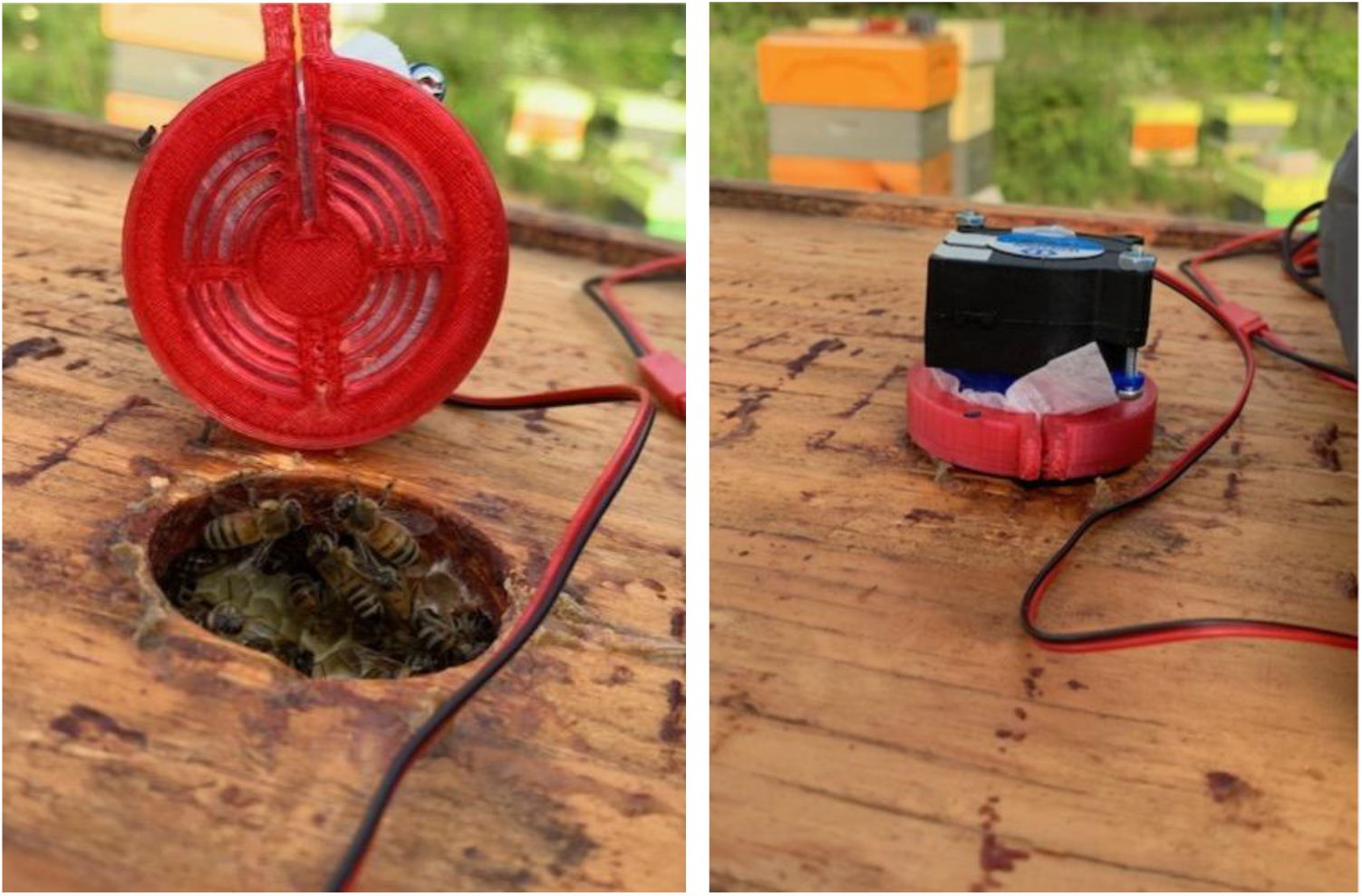
BeeSafe air sampler design. This is a modified version of the 12V mini sampler described by Garrett et al. 2023 with a new 3D printed top to prevent bees from landing on the filter surface.

### Study location

We collected air samples from honey bee hives from two apiaries, one near Fergus, Ontario, Canada and the other near Mono, Ontario, Canada (Figure 1). The collection sites geographical coordinates have been omitted to preserve the beekeeper’s identity. In total, 11 distinct beehive colonies were sampled, 5 from the apiary in Fergus and 6 in Mono. The Fergus hives were sampled once, whereas the Mono hives were sampled twice, in consecutive weeks, totaling 17 samples for this study (Table 1). Each air filter was left running for 5 to 6 hours and the filters were collected in a sterile ziplock bag. All honeybee colonies were inspected prior to sampling to ensure they were healthy (laying queen, eggs, brood) and free from obvious signs of disease.

**Table 1:** Sample collection ID organized by apiary, hive ID, collecting date, and air filtration sampling time.

| Sample_ID | Apiary | Hive_ID | Date | Time_sampling_air |
| --- | --- | --- | --- | --- |
| F01 | Fergus | PP4_141 | July 5<sup>th</sup> 2023 | 5h |
| F02 | Fergus | PP4_142 | July 5<sup>th</sup> 2023 | 5h |
| F03 | Fergus | PP4_143 | July 5<sup>th</sup> 2023 | 5h |
| F04 | Fergus | PP4_144 | July 5<sup>th</sup> 2023 | 5h |
| F05 | Fergus | PP4_145 | July 5<sup>th</sup> 2023 | 5h |
| M06 | Mono | S1B | May 31<sup>st</sup> 2023 | 6h |
| M07 | Mono | S2A | May 31<sup>st</sup> 2023 | 6h |
| M08 | Mono | S3A | May 31<sup>st</sup> 2023 | 6h |
| M09 | Mono | S4A | May 31<sup>st</sup> 2023 | 6h |
| M10 | Mono | S5A | May 31<sup>st</sup> 2023 | 6h |
| M11 | Mono | S5B | May 31<sup>st</sup> 2023 | 6h |
| M12 | Mono | S1B | June 08<sup>th</sup> 2023 | 6h |
| M13 | Mono | S2A | June 08<sup>th</sup> 2023 | 6h |
| M14 | Mono | S3A | June 08<sup>th</sup> 2023 | 6h |
| M15 | Mono | S4A | June 08<sup>th</sup> 2023 | 6h |
| M16 | Mono | S5A | June 08<sup>th</sup> 2023 | 6h |
| M17 | Mono | S5B | June 08<sup>th</sup> 2023 | 6h |

### DNA extraction, PCR, Sequencing

To extract DNA, we followed the methods described by Garrett et al. (2023) for airborne eDNA. In brief, from the square filters, a half circle was first cut from the filter's center using scissors and forceps, sterilizing them with 10% commercial bleach and 10% ethanol between each filter. The filter was then soaked in 3-4 ml of PBS and shaken in an incubator at 50-56°C overnight. The PBS was transferred into a microcentrifuge tube, spun at 6000 x g for 3 minutes, and the PBS was removed, leaving the pellet. This process was repeated until all PBS was used. The pellet was then processed using a Qiagen Blood and Tissue kit (Qiagen, Canada) with the following modifications: incubating for 15 minutes in step one, warming the elution buffer before use, incubating at room temperature for 5 minutes before elution, and eluted using only 100 µl of buffer.

PCR preparation was conducted inside an AirClean PCR cabinet situated in an isolated room to minimize contamination from other airborne sources. The cabinet was cleaned with a 50% commercial bleach/water solution and UV sterilized. The PCR protocols followed those outlined by Garrett et al. (2023).

We amplified the hypervariable region V4 of the 16S rRNA gene using the primers 515F: 5'-GTGYCAGCMGCCGCGGTAA-3' and 806R: 5'-GGACTACNVGGGTWTCTAAT-3', which were modified to include Illumina adaptor regions (Comeau et al., 2011; Herlemann et al., 2011; Klindworth et al., 2013). We added 2 µl of extracted DNA to a PCR reaction containing 6.0 µl of sterile water, 6.0 µl of Qiagen PCR Master Mix (Qiagen), and 0.5 µl of 10 µM of forward and reverse primers. The cycling conditions were as follows: an initial denaturation at 94 °C for 3 min, followed by 35 cycles of 94 °C for 45 s, 52.5 °C for 60 s, and 72 °C for 90 s, with a final elongation step at 72 °C for 10 min. For the ITS gene, we used the primersITS-S2F: 5'-ATGCGATACTTGGTGTGAAT-3' and ITS4: TCCTCCGCTTATTGATATGC modified with Illumina adapter regions (Cheng *et al*., 2016). The PCR reaction mixtures were the same as described above for 16S. The cycling conditions were as follows: an initial denaturation at 94 °C for 4 min, followed by 34 cycles of 94 °C for 30 s, 55 °C for 40 s, and 72 °C for 60 s, with a final elongation step at 72 °C for 10 min.

In addition to the air filter samples, we included control samples to ensure data quality as follow: negative controls (no template), positive controls for plants (*Cannabis sativa*), and bacteria (ZymoBIOMICS Microbial Community DNA Standard). All PCR products, including these controls (positive, negative, and extraction blank), were visualized on a 1.5% agarose gel. The amplified samples were then sent to the Queen Mary University of London Genome Center for library preparation which included bead clean up and size selection by 0.9x Ampure Beads (Beckman Coulter, Brea, Ca, USA) quantification, quality control (QC) and normalization using Qubit (Invitrogen, Carlsbad, CA, USA) nucleotide quantification, and DNA D100 Tape station (Agilent) and each PCR product was then independently barcoded using unique indexes. Sequencing was performed on an Illumina MiSeq v3 2 × 300 cycle run (Illumina San Diego, Ca, USA). The resulting reads were demultiplexed on site and exported as FASTQ files for subsequent bioinformatic analysis. Data are available in the Genbank SRA (will be public at article acceptance).

### Data analysis

We processed raw reads using the Quantitative Insights into Microbial Ecology (QIIME2) v. 2019.10 pipeline (Bolyen et al. 2019). Briefly, we removed primers and sequencing adapters using cutadapt (Martin, 2011). Subsequently, raw reads underwent denoising and quality filtering with the DADA2 plugin, which identifies and filters chimeric sequences, calls amplicon sequence variants (ASVs), and generates a feature table of ASV counts and host metadata (Callahan et al. 2016). We used the representative ASV sequences as input for taxonomic classification using the Naïve Bayesian feature classifier on the SILVA reference database trained on the V4 515F/806R region of the 16S rRNA gene (Bokulich et al. 2018). For the ITS gene we used RESCRIPt (REference Sequence annotation and CuRatIon Pipeline) which is a python package and QIIME 2 plugin for formatting, managing, and manipulating sequence reference databases (Robeson et al. 2021). The plant database name used for the parameter *--p-query* was *txid33090[ORGN]*, and for fungi was *177353[BioProject]*. Finally, the Naïve Bayesian feature classifier was used as described above.

Potential contaminants were identified using the R package **decontam** (v.1.18.0; Davis et al. 2018), based on their increased prevalence in negative controls within the ASV feature table. To identify the core set of ASVs for each gene, we employed the `core` function from the **microbiome** R package (Lahti et al., microbiome R package. URL: http://microbiome.github.io). This function returns taxa that exceed specified prevalence and detection thresholds; in our analysis, we applied a prevalence threshold of 50%. Details of the negative controls are provided in Supplement 1.

### Statistical Analyses

To assess the variability of sequence data between samples, we calculated beta diversity, grouped according to Location, and Collection-date. The phylogeny-based weighted and unweighted UniFrac distances (Lozupone and Knight, 2005; Lozupone et al. 2007) were computed using the *phyloseq* package (Murdie and Holmes, 2013). Additionally, principal coordinate analysis (PCoA) plots were generated in R based on weighted (considering sequence abundance) and unweighted (presence/absence) UniFrac distance matrices.

To test for differences in beta diversity according to the factors and levels mentioned above, we employed the Permutational Multivariate Analysis of Variance (PERMANOVA), which is an analog of MANOVA for partitioning distance matrices across a multivariate data cloud (Anderson, 2001). PERMANOVA analyses were conducted using the *adonis2* function in the *vegan* (v.2.6-4) package (Oksanen et al. 2007). We also performed an analysis of multivariate homogeneity (PERMDISP) (Anderson, 2006) using the *betadisper* function to test whether the groups differed in their dispersion. We performed diversity analyses using the programming language R (R Core Team, 2022) v.4.2.1.

## Results

Air filters running for 5-6 hours successfully retrieved DNA from the air inside all sampled honey bee colonies. In total, we obtained 233,479 reads for 16S, 149,091 reads for ITS plants, and 775 for ITS fungi. After filtering out ASVs present in control samples, we retained 165,898 reads for 16S, 94,896 for ITS plants, and 775 for ITS fungi, representing 3,619 ASVs for 16S, 573 ASVs for ITS Plants, and 18 ASVs for ITS Fungi.

The number of ASVs per sample varied across datasets (Table 2). For bacterial 16S, the ASV count ranged from 186 to 514. ITS Plants yielded between 17 and 85 ASVs, while ITS Fungi showed a much narrower range, with ASV counts between 1 and 6.

**Table 2.** Number of Amplicon Sequence Variants found in each air filter sampled.

| Sample_ID | Apiary | ITS_Plant ASVs | ITS_Fungi ASVs | 16S_Bacterial ASVs |
| --- | --- | --- | --- | --- |
| F01 | Fergus | 85 | 6 | 235 |
| F02 | Fergus | 17 | 2 | 193 |
| F03 | Fergus | 30 | 4 | 168 |
| F04 | Fergus | 19 | 2 | 255 |
| F05 | Fergus | 19 | 2 | 227 |
| M06 | Mono | 24 | 1 | 248 |
| M07 | Mono | 66 | 1 | 243 |
| M08 | Mono | 60 | 3 | 267 |
| M09 | Mono | 42 | 2 | 233 |
| M10 | Mono | 50 | 1 | 238 |
| M11 | Mono | 28 | 1 | 186 |
| M12 | Mono | 51 | 1 | 498 |
| M13 | Mono | 27 | – | 359 |
| M14 | Mono | 56 | 1 | 374 |
| M15 | Mono | 59 | 2 | 514 |
| M16 | Mono | 76 | 1 | 340 |
| M17 | Mono | 73 | 1 | 373 |

### Microbial honey bee hive air samples diversity

Among the top 25 most frequent 16S ASVs, the microbial composition varied between the two apiaries. At Fergus apiary, across the five colonies (F01-F05), the dominant taxa were *Sphingomonas* sp. (22.8%), *Pseudomonas* sp. (20.2%), and *Massilia* sp. (19.8%). In contrast, at Mono apiary, across six colonies (M06-M17), the predominant ASVs in week 1 were *Bartonella* sp. (28.7%) and *Gilliamella* sp. (10.9%), shifting in week 2 to *Bartonella* sp. (22.1%) and *Nocardiopsis* sp. (18.4%) (Figure 2).

**Figure 2:**
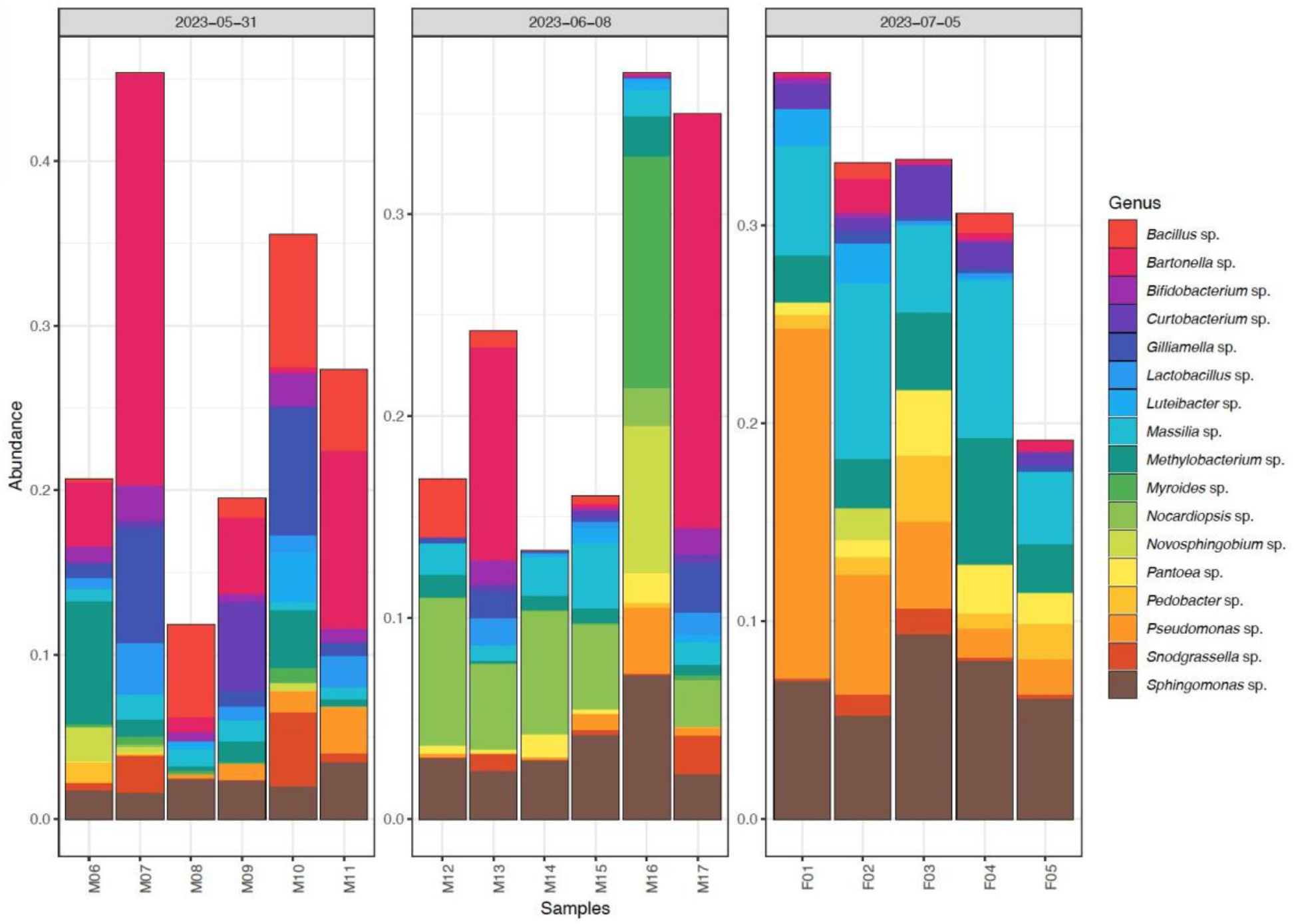
Microbiota composition of managed honey bee hives, illustrating the relative abundance of the top 25 bacterial genera. Each stacked bar corresponds to an individual air sample. Hives are grouped by location and time of collection. M06 to M17 represent six hives from the Mono apiary, sampled at two time points: M06 to M11 were collected on May 31, 2023 (week 1), and M12 to M17 on June 6, 2023 (week 2). In both sampling weeks, the microbiota was primarily dominated by *Bartonella* sp. F01 to F05 represent hives from the Fergus apiary, where the microbiota was predominantly composed of *Sphingomonas* sp.

The core set (present in at least 50% of samples) of ASVs identified from the 17 honey bee air samples included 32 ASVs. Among these, the ASV with the highest prevalence, detected in 100% of the samples and accounting for 2.59% of the relative abundance, belonged to the Phylum *Proteobacteria* (Family Sphingomonadaceae, Genus *Sphingomonas*). Additionally, the ASV with the highest relative abundance, comprising 5.06% of the total, was found in 88% of the samples and was classified within the Phylum *Proteobacteria* (Family Rhizobiaceae, Genus *Bartonella*).

### Air DNA captures honey bee foraging

Among the top 25 most frequent ITS ASVs for plants, *Brassica* sp. dominated the data, accounting for 49% across the five colonies at the Fergus apiary. At the Mono apiary, during week 1, *Brassica* sp. remained predominant (46%), followed by *Rhamnus* sp. (24.1%). In week 2 at the Mono apiary, the plant ASVs were primarily *Hieracium* sp. (30.2%) and *Pilosella* sp. (28.8%) (Figure 3).

**Figure 3:**
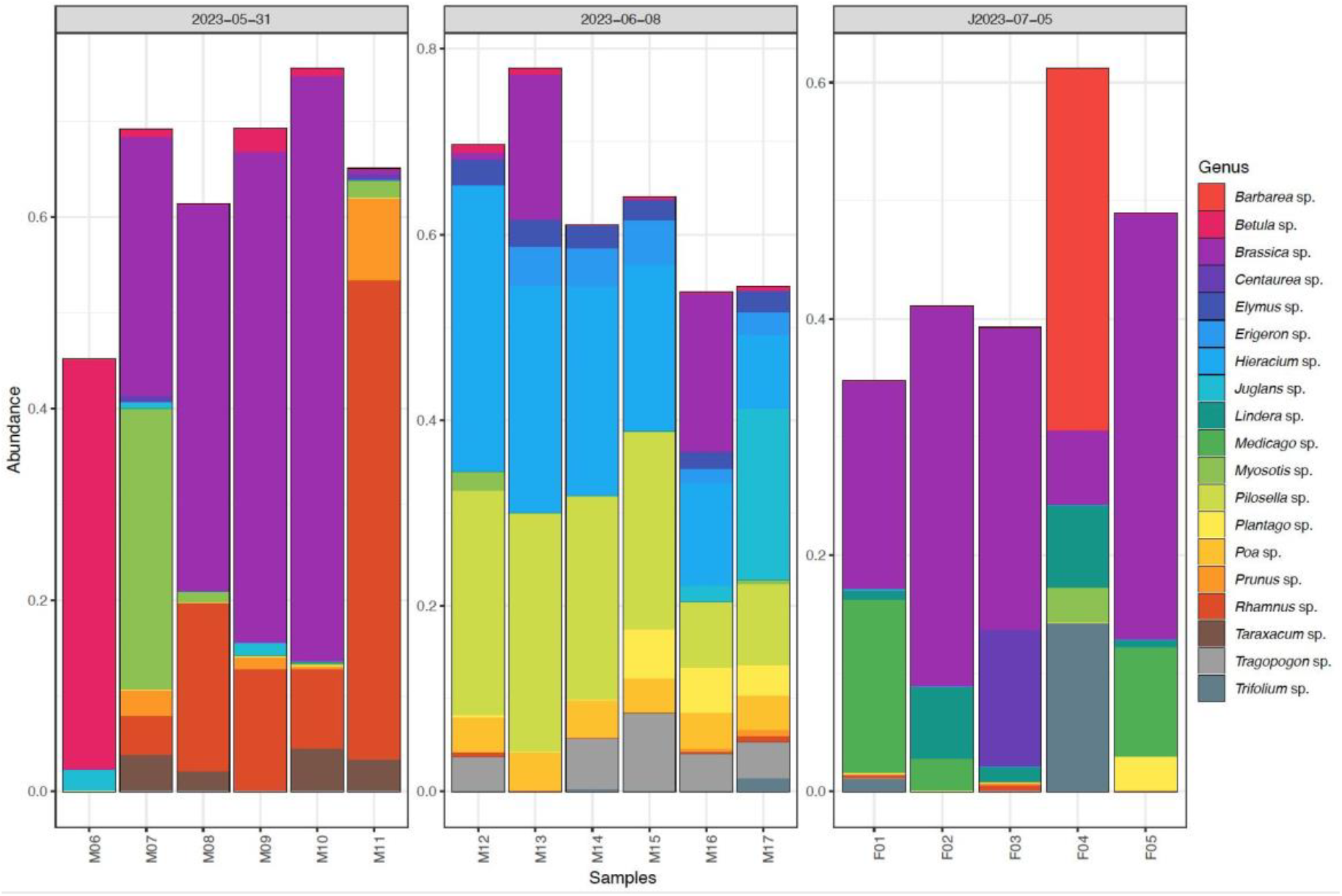
ITS plant composition of managed honey bee hives, showing the relative abundance of the top 25 plant genera. Each stacked bar represents an individual air sample. Hives are grouped by location and time of collection. M06-M17 represent six hives from the Mono apiary sampled twice: M06-M11 were collected on May 31, 2023 (week 1), and M12-M17 on June 06, 2023 (week 2). Week 1 hives were dominated by *Brassica* sp. and *Rhamnus* sp. Week 2 was dominated by *Hieracium* sp. and *Pilosella* sp. F01 to F05 are samples from hives at the Fergus apiary and collected July 5, 2023 (week 3), where the plant composition is dominated by *Brassica* sp.

The core set of ASVs across the 17 honey bee air samples consisted of 2 ASVs. The ASV with the highest prevalence (70.59%) and a relative abundance of 5.50% belonged to the Phylum Streptophyta (Family Brassicaceae, Genus *Brassica*). Additionally, the ASV with the highest relative abundance (8.30%) and a prevalence of 41.18% also belonged to Streptophyta (Family Brassicaceae, Genus *Brassica*).

### Plant and bacterial community beta diversity

Beta diversity was evaluated to assess the impact of apiary sites and sampling dates on the composition of ITS and 16S in the samples. Cluster analysis, both weighted and unweighted, revealed that the samples grouped by sampling date and apiary site (Figure 4), with greater variability in sequence recovery between sites than within them. The weighted analysis of ITS Plant diversity explained 66.7% of the variance (Axis 1 and 2), with samples clustering according to both sampling date and apiary location. Similarly, the weighted analysis of 16S microbial diversity explained 44.3% of the variability (Axis 1 and 2).

**Figure 4:**
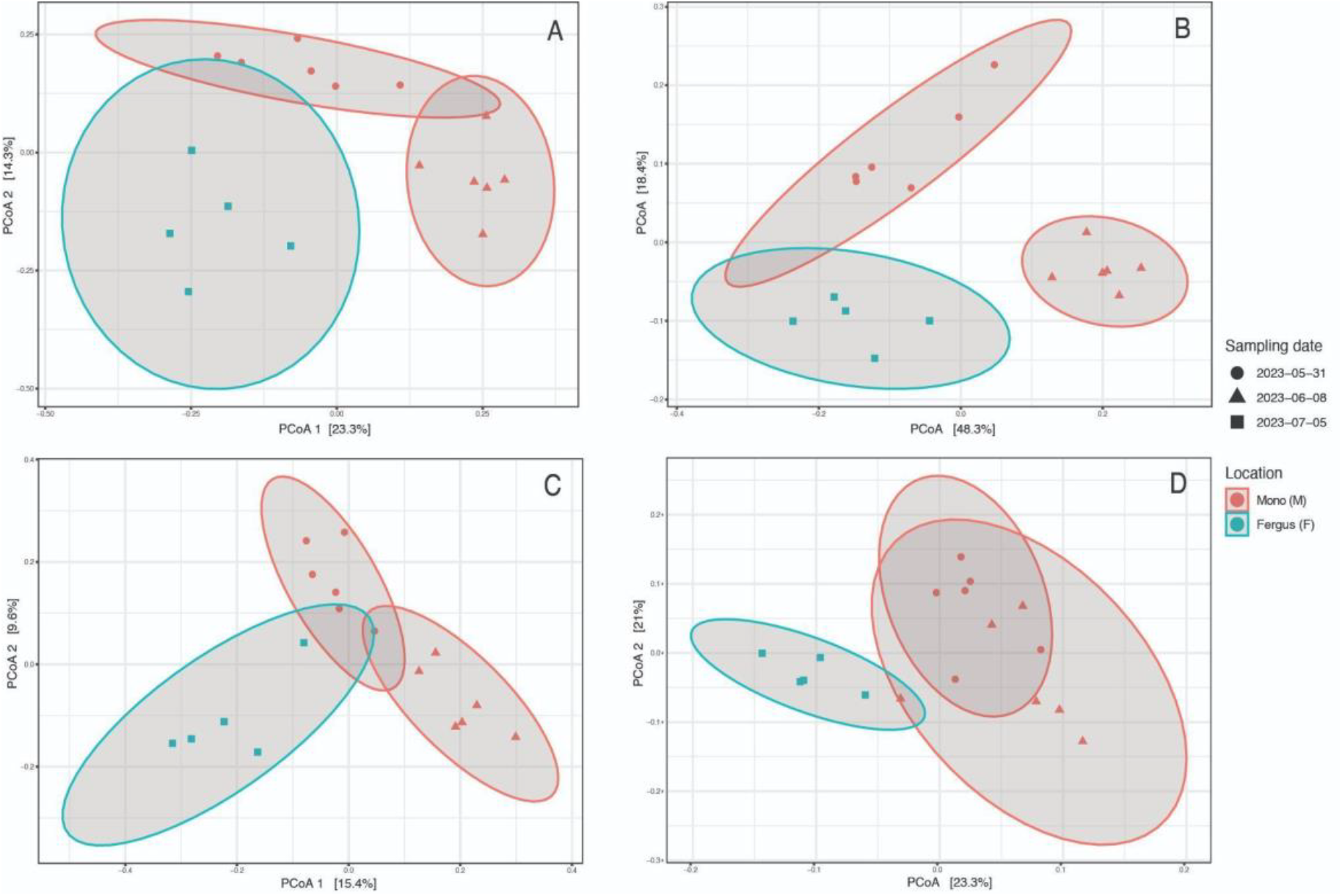
Principal Coordinate Analysis (PCoA) plots of plants (A, B) and microbial (C, D) communities, unweighted (A, C) and weighted (B, D) UniFrac distances of apiary location (Mono and Fergus) and date collected. Unweighted considers presence/absence data, whereas weighted considers read abundance in the composition. A higher percentage of data variability on Axis 1 is explained by composition/ abundances (Weighted UniFrac) in both ITS Plants and 16S Microbial data.

### 16S Microbial

The PERMANOVA weighted UniFrac analysis indicated that location accounted for the largest portion of data variation (adonis2, 20.2%, F = 4.1, p = 0.0001), followed by sampling date, which explained 10.9% of the variation (adonis2, F = 2.2, p = 0.0150). Similarly, the PERMANOVA unweighted UniFrac analysis showed that location explained the greatest amount of variation (adonis2, 12.3%, F = 2.2, p = 1e-04), with sampling date accounting for 10.8% of the variation (adonis2, F = 2.0, p = 3e-04).

However, the results of the PERMANOVA analysis could be affected by non-homogeneous dispersion of the data in the weighted UniFrac analysis. This can be seen in the location variable (betadisper, F = 5.30, P < 0.04), but not for the variable sampling date (betadisper, F = 1.42, P = 0.28). For the unweighted UniFrac, all variables are homocedastic.

### ITS Plants

The PERMANOVA weighted UniFrac analysis suggests that sampling date accounted for the largest portion of data variation (adonis2, 32.2%, F = 10.9, p = 1e-04), followed by location, which explained 26.3% of the variation (adonis2, F = 8.9, p = 1e-04). In the unweighted UniFrac analysis, location was the primary contributor to data variation (adonis2, 17.2%, F = 3.6, p = 2e-04), while sampling date explained 16.4% (adonis2, F = 3.4, p = 1e-04). Both weighted and unweighted UniFrac variables were found to be homoscedastic.

### ITS Fungi honey bee hives air samples diversity

ITS fungi diversity was much smaller than ITS plants and bacteria, with a total of 18 ASVs. Given that we obtained a small number of reads and ASVs for ITS Fungi, we provide only the histogram, without any further analysis (Figure 5). These data were dominated by *Alternaria*, a ubiquitous fungus, found in soil, air, and plant debris, that thrive in warm, humid conditions (Woundenberg et al., 2013), extremely common in eDNA metabarcoding results (Clare, personal observation).

**Figure 5:**
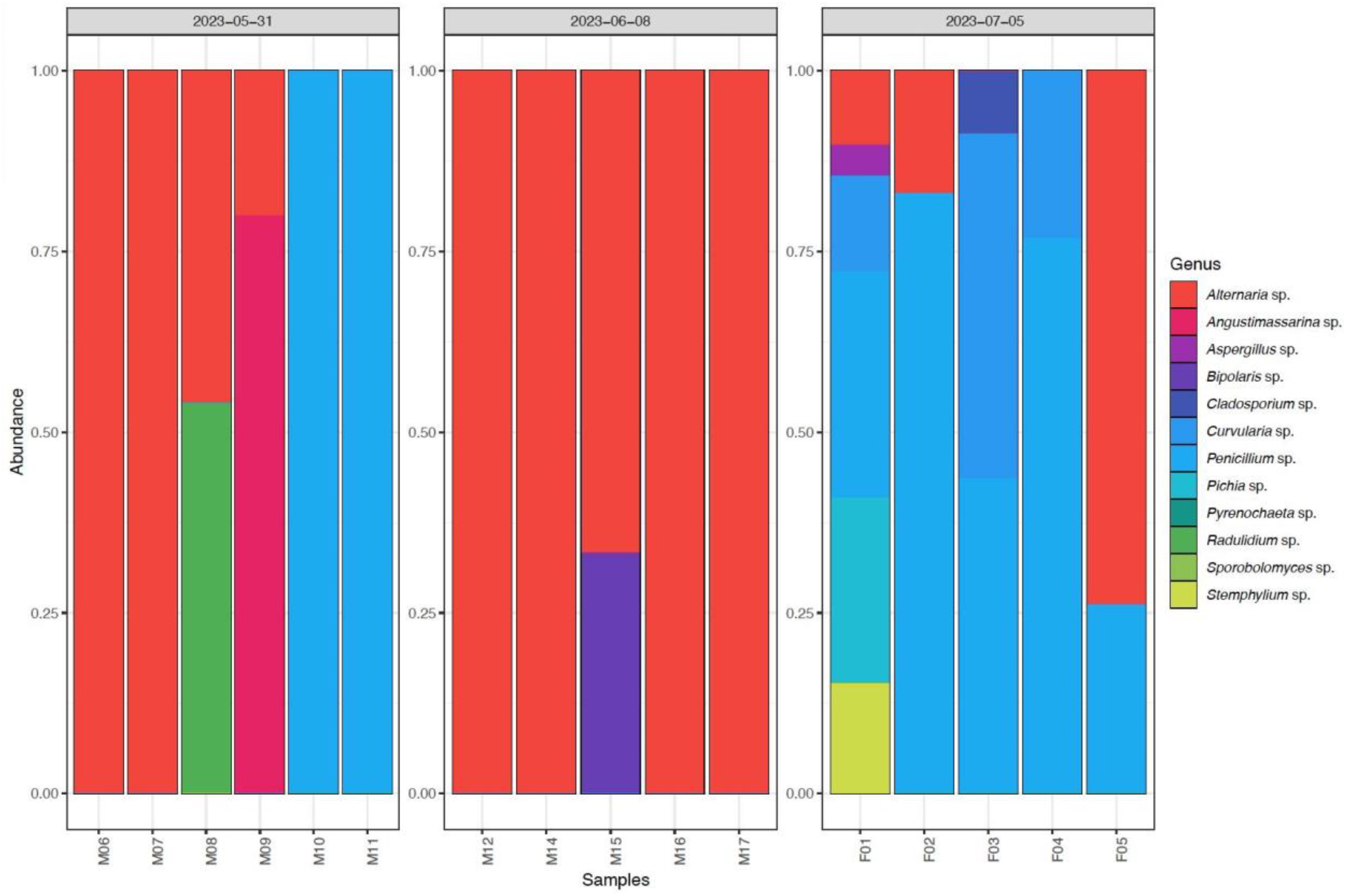
ITS fungi composition of managed honey bee hives, showing the relative abundance of the top 25 fungi genera. Each stacked bar represents an individual air sample. Hives are separated by location and time of collection. M06-M17 represent six hives from the Mono apiary sampled twice: M06-M11 were collected on May 31, 2023 (week 1), and M12-M17 on June 06, 2023 (week 2). Week 1 hives were dominated by *Alternaria* sp. and *Penicillum* sp, whereas Week 2 was dominated by *Alternaria* sp. Alternaria is extremely common in airborne eDNA work and should be considered nearly universal. F01 to F05 are hives from the Fergus apiary collected July 5, 2023, with fungi composition dominated by *Penicillium* sp.

## Discussion

The increasing use of hive sharing and rental hives for both personal use and large-scale agricultural pollination services means rapid diagnostics are vital. This includes assessing colony health in terms of foraging, microbial health and pathogen detection. In many cases, rapid early detection of problems or screening for hive activities prevents larger losses, and these screenings are sometimes mandated by regulatory entities to safeguard the apiculture community. In this study we report on a pilot study in non-invasive air sampling for diagnostics of foraging and microbiome analysis. Our data suggest that this may be a very effective tool for rapid ecological assessment of colonial bees and provides an effective way to monitor recent activity patterns at the colony level.

### Monitoring microbiome for honey bee health

Honey bees have a relatively simple gut microbiome consisting of a core of five well characterized bacterial lineages (Motta and Moran 2024). This core of bacterial species provides a protective function to individuals helping in metabolism (Zheng et al. 2016), protection from harmful pathogens (Raymann and Moran 2018), digestion (Engel et al. 2012) and detoxification (Zheng et al. 2016). An intact microbiome is also thought to be required for effective gene expression linked to development and behaviour (Zhang et al. 2022), though these helpful microbes can be disrupted by pesticides. The main core bacteria includes taxa within the *Bombilactobacillus, Bifidobacterium, Gilliamella, Lactobacillus* and *Snodgrassella* (Motta and Moran 2024). Many other pathogenic bacteria and fungi have been associated with *Apis* and some, like *Paenibacillus* which causes foolbroud, are deadly and in many jurisdictions must be reported to authorities when detected. Monitoring microbial health of honey bees is thus a commercial and regulatory requirement. Given the importance of this type of monitoring it was interesting that even in this pilot study, the honey bee core gut microbiome was retrieved in most of the samples. From the 17 colonies analyzed, *Lactobacillus melliventris* (Olofsson TC et al. 2014) was sampled in 10; *Bifidobacterium asteroids* (Milani et al., 2014) in 13; *Snodgrassella alvi* in 13; *Gilliamella apicola* in 15; and *Bartonella apis* in 15. In those same colonies we detected no known pathogenic taxa, a result that is almost certainly accurate (no false negatives) as these colonies are well monitored and had been screened for these using traditional approaches. Using airborne eDNA for monitoring honey bee colony health shows promise and accurately reflected what was known of these colonies. However, a directly compared airborne eDNA results to more traditional methods of gut microbiome analysis, would be a logical next step to quantify the effectiveness of this recovery method.

### eDNA and plant-bee interactions

There are a variety of approaches to study pollination using DNA or eDNA techniques. The choice of method likely depends on the question being asked and each approach has its own advantages and disadvantages. Important questions include whether the objective is to study the behavior of the insect or the function of pollination and whether we are interested in individuals or communities. For example, here we are interested primarily in the *Apis* colony as a whole and so measurements are made at the colony level. In contrast, if our objective was to study pollination we might prefer to swab plants for animal eDNA (Newton et al. 2023) to gain a broader picture of animal visitation. Similarly, if we are interested in individual movements, or home ranges, measurements at the level of individual workers (e.g. pollen metabarcoding Bell et al. 2017) would be more advantageous. One advantage of colony level measurements is the presumably lower risk of false positives detections i.e. we can assume most plant DNA inside was brought in by the bees. This contrasts with measurements of insect DNA on leaves where some or all might be from the environment rather than a specific interaction (Valentin et al. 2018). While the exact definition of eDNA is variable and may refer specifically to extra-cellular DNA or more generally to any sample collected from an environment, we suspect a substantial part of what is being analyzed here is full cells (e.g. microbes) and at least some may be multicellular debris including pollen. It would be impossible (and meaningless) to attempt to distinguish the samples in this way, but we might expect more intact material to generate larger DNA concentrations and thus contribute to more of the eventual data being analyzed. While we treat these collectively as an environmental sample, we do point out that only some constitutes eDNA in the strict sense.

Two interesting challenges in using this method will be to first, quantify the interaction between hive air and honey as a DNA source to truly separate recent form past temporal signals, and second, develop effective negative controls. While the first might be accomplished by honey metabarcoding in parallel to air sampling, developing negative field controls in airborne sampling is exceedingly complex. In aquatic eDNA, sterile water can be run through equipment to gauge potential field contaminants; however, no real equivalent exists for air. Several proposed options are only partially effective. For example, deploying unused/unpowered samplers may control for the suction effect but fail to account for air current drift, and “contaminants” may in fact be real signals (Clare pers observation). Similarly using leaves to control for environmental accumulation on flower petals when tracking insect visitation is likely over cautious because the same insects visiting flowers likely generate DNA in a nearby air column. Negative controls within colonies are equally complex. Like the leaves on the plant, the honey itself is likely too conservative since common plant DNA sources would be in the air and in the honey. An air filter outside could generate a list of what “could” be simple environmental accumulation, but the same things are also likely foraging targets and exclusion would be overly conservative. One advantage of air over surfaces (like flowers) is that the DNA appears to settle quickly (Garrett et al. 2023). While we expect DNA to settle out of the air and onto substrates (flowers, leaves, soils) and thus build up, air appears to be a much more dynamic DNA source. Creativity in developing negative field controls will be required.

### The potential of non-invasive monitoring

There may be many advantages to ongoing or hive health monitoring from eDNA that are not found in traditional approaches. eDNA is often cited as being surprisingly good at detecting rare taxa and may identify an infection or presence of mites before it becomes established, something considered key in controlling outbreaks (Thomsen & Willerslev, 2015). On the other hand, caution needs to be taken when using this as a tool in hive health assessments. Many microbial, viral and parasitic diseases of bee hives require official reporting or even hive destruction (e.g. foulbrood). While air-based monitoring could prove an effective method to accelerate early detection, confirmation by follow up inspection is required to mitigate any false positives. Since airborne eDNA sampling is in its infancy and the duration and travel distance of such signals are still under active investigation, we recommend any health related data be considered an early warning only and backed up with conventional sampling under local inspection laws.

However, the potential for new ecological investigation is substantial. Our observations suggest a high level of variability in the recovery of taxa between samples, and while we do not treat read counts as ecologically meaningful, they demonstrate high variability between samples, dates and sites. This provides a strong argument in favour of increased replication to understand the spatial temporal signature of the samples. The ability to isolate and monitor recent hive foraging through colony air opens up a number of new ecological research opportunities. If eDNA signals in air are of short duration, fine scale monitoring of hive activity is possible without actually touching individuals and multiple hives can be monitored simultaneously without field labour. Monitoring daily, or even hourly patterns of foraging becomes an option in a way not possible with alternative approaches. Questions about fine scale responses to weather and environmental perturbations would be easier to quantify with eDNA, and the effect of experimental manipulation of resources could be quantified rapidly at the full hive community level.

## Acknowledgements

This work was funded by the Natural Sciences and Engineering Research Council of Canada through the Discovery Grants Program, (*RGPIN-2021-03611*), the Government of Canada’s New Frontiers in Research Fund NFRF-E program (*NFRFE-2022-00277)* to ELC and AZ, and NFRF-T program (NFRFT-2020-0073) Tracing the Patterns of Life on a Changing Planet, Genome Canada (BIOSCAN Canada) and Ontario Genomics (OGI-208).

